# Synergistic interplay between PHF8 and HER2 signaling contributes to breast cancer development and drug resistance

**DOI:** 10.1101/682476

**Authors:** Qi Liu, Nicholas Borcherding, Peng Shao, Peterson Kariuki Maina, Weizhou Zhang, Hank Heng Qi

## Abstract

HER2 plays a critical role in tumorigenesis and is associated with poor prognosis of HER2-positive breast cancers. Although, anti-HER2 drugs show benefits in breast cancer therapy, *de novo* or acquired resistance often develop. Epigenetic factors have been increasingly targeted for therapeutic purposes, however, such mechanisms interacting with HER2 signaling are poorly understood. This study reports the synergistic interplay between histone demethylase PHF8 and HER2 signaling, i.e. PHF8 is elevated in HER2-positive breast cancers and is upregulated by HER2; PHF8 plays coactivator roles in regulating *HER2* expression and HER2-driven epithelial-to-mesenchymal transition (EMT) markers and cytokines. The HER2-PHF8-IL-6 regulatory axis was proved both in cell lines and in the newly established *MMTV-Her2/MMTV-Cre/Phf8^flox/flox^* models, with which the oncogenic function of Phf8 in breast cancer *in vivo* was revealed for the first time. Furthermore, PHF8-IL-6 axis contributes to the resistance of Trastuzumab *in vitro* and may play a critical role in the infiltration of T-cells in HER2-driven breast cancers. This study reveals novel epigenetic mechanisms underlying HER2-driven cancer development and anti-HER2 drug resistance.

## Introduction

Breast cancer is the most commonly diagnosed cancer and is the second leading cause of cancer death in American women. About 268,600 new cases of breast cancer will be diagnosed and about 41,760 women will die from breast cancer in 2019 in the United States (Siegel et al, 2019). Breast cancers are grouped into three categories, which are not mutually exclusive: ER (estrogen receptor)-positive; ERBB2/HER2/NEU (HER2 is used hereafter)-positive (HER2+) and triple negative. HER2+ breast cancers occur in 20-30% of breast cancer and are often associated with poor prognosis (Roskoski, 2014). HER2 is a transmembrane receptor tyrosine kinase and plays critical roles in the development of both cancer and resistance to therapy in the cases of both HER2+ (Baselga & Swain, 2009; Roskoski, 2014) and HER2-negative (HER2-) (Cao et al, 2009; Duru et al, 2012; Hurtado et al, 2008) breast cancers. In the later cases, such as luminal or triple-negative breast cancer, HER2 expression is elevated within a defined group of cancer stem cells that are believed to be the true oncogenic population in the heterogeneous breast cancer and to confer resistance to both hormone and radiation therapies (Cao et al, 2009; Duru et al, 2012; Hurtado et al, 2008). Trastuzumab, a humanized anti-HER2 antibody and Lapatinib, a HER2 kinase inhibitor, dramatically improved the treatment of HER2+ breast cancer and gastric cancer patients (Iqbal & Iqbal, 2014). Notably, these anti-HER2 therapies also exhibited a benefit to HER2-cancer patients (Paik et al, 2008). However, drug resistance often develops *de novo* and becomes another obstacle in successful therapy (Roskoski, 2014). Thus, to identify novel therapeutic targets that are critical for HER2-driving tumor development and resistance to therapy is still needed.

The importance of epigenetic mechanisms in cancer development has been recognized and chromatin regulators have been increasingly targeted in developing cancer therapies (Greer & Shi, 2012; Verma & Banerjee, 2015). For example, targeting of the bromodomain and extra terminal domain (BET) protein by the inhibitor JQ1 has been shown to antagonize the proliferation of multiple myeloma cells through repressing c-MYC and its downstream effectors (Delmore et al, 2011). Similarly, targeting the histone demethylase KDM4 family member, NCDM-32B, has been effective in reducing the proliferation and transformation of breast cancer cells (Ye et al, 2015). In context of HER2, the association of epigenetic changes including DNA methylation, histone modifications, and ncRNAs/miRNAs with HER2+ breast cancer susceptibility was critically reviewed (Singla et al, 2017). Importantly, histone deacetylase (HDAC) and DNA methylation inhibitors can upregulate *HER2* expression (Ramadan et al, 2018; Singla et al, 2017). Moreover, methylations on histone 3 lysine 4 (H3K4me3) and histone 3 lysine 9 (H3K9me2) are associated with the activation and downregulation of *HER2*, respectively (Singla et al, 2017). In fact, WDR5, a core component of H3K4me3 methyltransferase and G9a, the H3K9me2 methyltransferase, were claimed to be responsible for the changes of these modifications (Singla et al, 2017). However, whether and how histone demethylase, another major contributor to the epigenetic mechanisms, to *HER2* expression and HER2-driven tumor development and resistance to therapy remain largely unknown.

We have recently reported that histone demethylase PHF8 (PhD finger protein 8) promotes epithelial-to-mesenchymal transition (EMT) and contributes to breast tumorigenesis (Shao et al, 2017). We also demonstrated that higher expression of PHF8 in HER2+ breast cancer cell lines and functional requirement of PHF8 for the anchorage-independent growth of these cells. The demethylase activities of PHF8 were simultaneously identified towards several histone substrates: H3K9me2, H3K27me2 (Feng et al, 2010; Fortschegger et al, 2010; Kleine-Kohlbrecher et al, 2010; Loenarz et al, 2010) and H4K20me1 (Liu et al, 2010; Qi et al, 2010). These studies also revealed a general transcriptional coactivator function of PHF8. The following studies demonstrated the overexpression and oncogenic functions of PHF8 in various types of cancers such as prostate cancer (Bjorkman et al, 2012; Maina et al, 2016), esophageal squamous cell carcinoma (Sun et al, 2013), lung cancer (Shen et al, 2014), and hepatocellular carcinoma (Zhou et al, 2018). Beyond overexpression, the post-transcriptional and post-translational regulations of PHF8 were also recently elucidated. We identified the c-MYC-miR-22-PHF8 regulatory axis, through which the elevated c-MYC can indirectly upregulates *PHF8* by represses *miR-22*, a microRNA that targets and represses *PHF8* (Maina et al, 2016; Shao et al, 2017). Moreover, USP7-PHF8 positive feedback loop was discovered, *i.e*. deubiquitinase USP7 stabilizes PHF8 and PHF8 transcriptionally upregulates USP7 in breast cancer cells (Wang et al, 2016). With such mechanism, the stabilized PHF8 upregulates its target gene *CCNA2* to augment breast cancer cell proliferation. All these data support the elevated expression of PHF8 in cancers and its oncogenic roles. However, the epigenetic regulatory role of PHF8 plays in HER2-driving tumor development and resistance to anti-HER2 therapy remains largely unknown.

In this study, we report the elevation of PHF8 in HER2+ breast cancers and by HER2 overexpression. The upregulated PHF8 plays a coactivator role in *HER2* expression and genes that upregulated by activated HER2 signaling. Moreover, we revealed that PHF8 facilitates the upregulation of IL-6 both *in vitro* and *in vivo* and the PHF8-IL-6 axis contributes to the resistance to anti-HER2 drugs. This study sheds a light on the potential of drug development on inhibition of histone demethylase in HER2-driven tumor development and therapy resistance.

## Results

### PHF8 expression is elevated in HER2+ breast cancers and is upregulated by HER2

Prompted by our previous findings of higher expression of PHF8 in HER2+ breast cancer cells and its functional requirement in the anchorage-independent growth of these cells (Shao et al, 2017), we first evaluated *PHF8* mRNA levels in breast cancers with recent RNA-sequencing (RNA-seq) data through Gene Expression Profiling Interactive Analysis (GEPIA) (Tang et al, 2017). Intriguingly, *PHF8* mRNA levels are only slightly elevated across breast cancers (n=1085) as well as in subtypes of breast cancers, compared with normal tissue (n=291) (Supplemental Figure 1A and B). As PHF8 is subject to both post-transcriptional and post-translational regulations such as c-MYC-miR-22-PHF8 (Maina et al, 2016; Shao et al, 2017) and USP7-PHF8 (Wang et al, 2016) regulatory mechanisms, the actual PHF8 protein levels in cancers can differ from its mRNA levels. We increased our breast cancer samples from previous study to a pool of samples of 486 breast cancer, 20 normal breast tissues, and 50 metastatic lymph nodes (US BIOMAX). PHF8 immunohistochemistry (IHC) staining on these breast cancer tissue arrays demonstrates significant elevation of PHF8 protein levels (strong nuclear staining) across breast cancer and in all subtypes, including HER2+ breast cancers (Supplemental Table 1 and Figure 1A). These data, together with higher PHF8 protein levels in HER2+ breast cancer SKBR3 and BT474 cells (Shao et al, 2017), suggest potential regulation of PHF8 by HER2. Continuing with this hypothesis, we found the overexpression of HER2 by pOZ retroviral system upregulates PHF8 at both protein and mRNA levels in MCF10A and MCF7 cells (Figure 1B). Conversely, *HER2* knockdown with siRNAs against *HER2* gene body compromised the upregulation of PHF8 proteins in these cells, but, rescued *PHF8* mRNAs dominantly in MCF7 cells (Figure 1C). *HER2* knockdown in SKBR3 and BT474 cells also downregulated PHF8 protein levels (Supplemental Figure 2). Moreover, a significant positive correlation between *HER2* and *PHF8* mRNAs is observed through the analysis of the Cancer Genome Atlas Breast Invasive Carcinoma (TCGA-BRCA), TCGA normal breast tissue and Genotype-Tissue Expression (GTEx) Program mammary tissue (Figure 1D). Taken together, these findings support our hypothesis that elevated HER2 upregulates PHF8.

**Figure 1.**
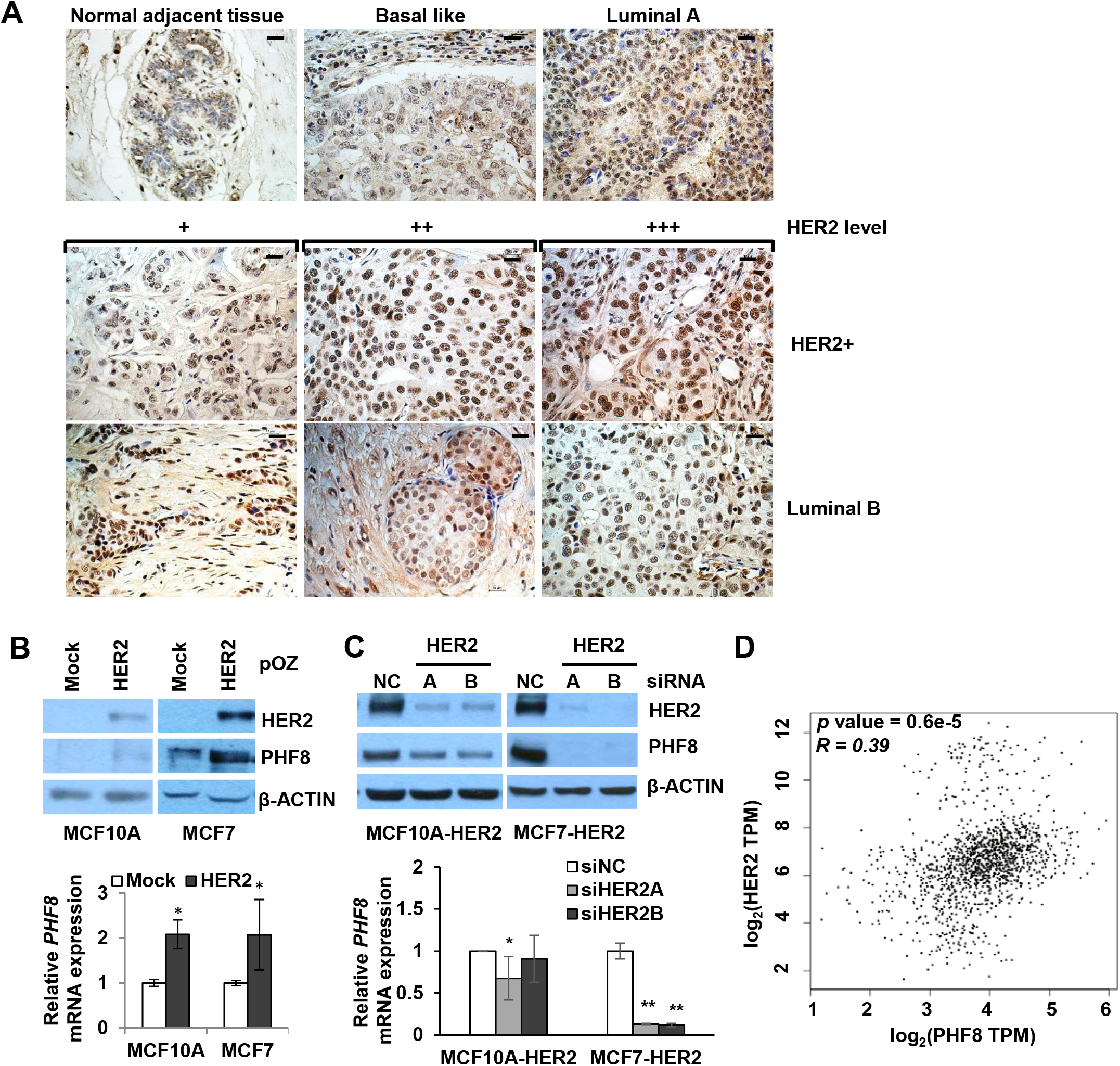
PHF8 expression is elevated in HER2+ breast cancers and is upregulated by HER2. **A**. Immunohistochemistry (IHC) of PHF8 in breast cancer tissue arrays. Representative PHF8 staining in cancer adjacent normal adjacent breast tissue, Basal like, Luminal A, Luminal B and HER2+ samples. Magnification, ×200, Bar: 10 μm. **B**. In MCF10A and MCF7 cells with or without HER2 overexpression, PHF8 protein (upper panel) and mRNA (lower panel) levels were assessed by western blotting and RT-PCR, respectively. **C**. In MCF10A-HER2 and MCF7-HER2 cells with or without HER2 knockdown, PHF8 protein (upper panel) and mRNA (lower panel) levels were assessed by western blotting and RT-PCR, respectively. All quantitative data are expressed relative to the value for control cells and are means ± SD from three independent experiments. \**p*<=0.05; **: *p*<=0.01. **D**. Scatter plot of *PHF8* and *HER2* expression levels from TCGA-BRCA tumor dataset, normal dataset and Genotype-Tissue Expression (GTEx) Program mammary tissue datasets.

### PHF8 is a transcriptional coactivator of *HER2* gene

Due to the coactivator role of PHF8, we hypothesize that PHF8 participates in the transcriptional regulation of *HER2*. PHF8 ChIP-seq data (Chromatin immunoprecipitation following by deep sequencing) from human embryonic H1 cells and K562 cells (Ram et al, 2011) show enrichments of PHF8 on the two promoter regions of *HER2* (Supplemental Figure 3). It is not surprising that PHF8 is co-localized with H3K4me3 as PHF8 binds to H3K4me3 via its PHD domain (Qi et al, 2010). Importantly, we identified similar enrichments of PHF8 on *HER2* promoters in SKBR3, BT474, and HCC1954 cells (Figure 2A). Loss-of-function of *PHF8* by either siRNA or shRNAs in SKBR3, BT474 and HCC1954 cells uniformly downregulated HER2 at both protein and mRNA levels (Figure 2B), supporting the coactivator function of PHF8 in the transcriptional regulation of *HER2*. We have recently identified a novel HER2 gene body enhancer (HGE), which recruits transcription factor TFAP2C (Liu et al, 2018a), a known positive regulator of *HER2* gene (Ailan et al, 2009; Bosher et al, 1996; Kulak et al, 2013; Perissi et al, 2000; Vernimmen et al, 2003). Interestingly, our early work showed that *TFAP2C* is a direct target gene of PHF8 (Qi et al, 2010). Thus, we ask if PHF8 regulates TFAP2C in HER2+ cells. *PHF8* knockdown downregulates TFAP2C protein and mRNA levels (Figure 2B and Supplemental Figure 4). Consequentially, the enrichments of TFAP2C at *HER2* promoters are reduced in the PHF8-RNAi cells (Figure 2C), implicating that the regulation of TFAP2C by PHF8 could contribute to *HER2* expression. *HER2* gene is constitutively active in these HER2+ breast cancer cells, rendering active chromatin status, therefore, the PHF8 demethylation substrates such as H3K9me2, H4K20me1 and H3K27me2 may not be highly enriched at *HER2* promoters. In contrast, H3K4me3 is known to be critical for the transcriptional regulation of *HER2* in breast cancer cells (Mungamuri et al, 2013). Importantly, our early studies showed that PHF8 also plays a critical role to sustain the level of H3K4me3 in various cell types (Maina et al, 2017; Qi et al, 2010). ChIP experiments re-enforced this role of PHF8: *PHF8* knockdown reduced the H3K4me3 levels at *HER2* promoters in all three cell lines tested (Figure 2C). These data support that PHF8 participates in the transcriptional regulation of *HER2* by sustaining H3K4me3 levels. Moreover, H3K27ac, a general activation marker, is downregulated on *HER2* promoter regions in *PHF8* knockdown cells (Figure 2C). In this context, PHF8 may execute its demethylation activity on H3K27 to prime the acetylation or indirectly regulate H3K27ac through its acetyltransferase. Taken together, we revealed the role of PHF8 in the transcriptional regulation of *HER2*, the underlying mechanisms may involve direct and indirect regulations of multiple factors.

**Figure 2.**
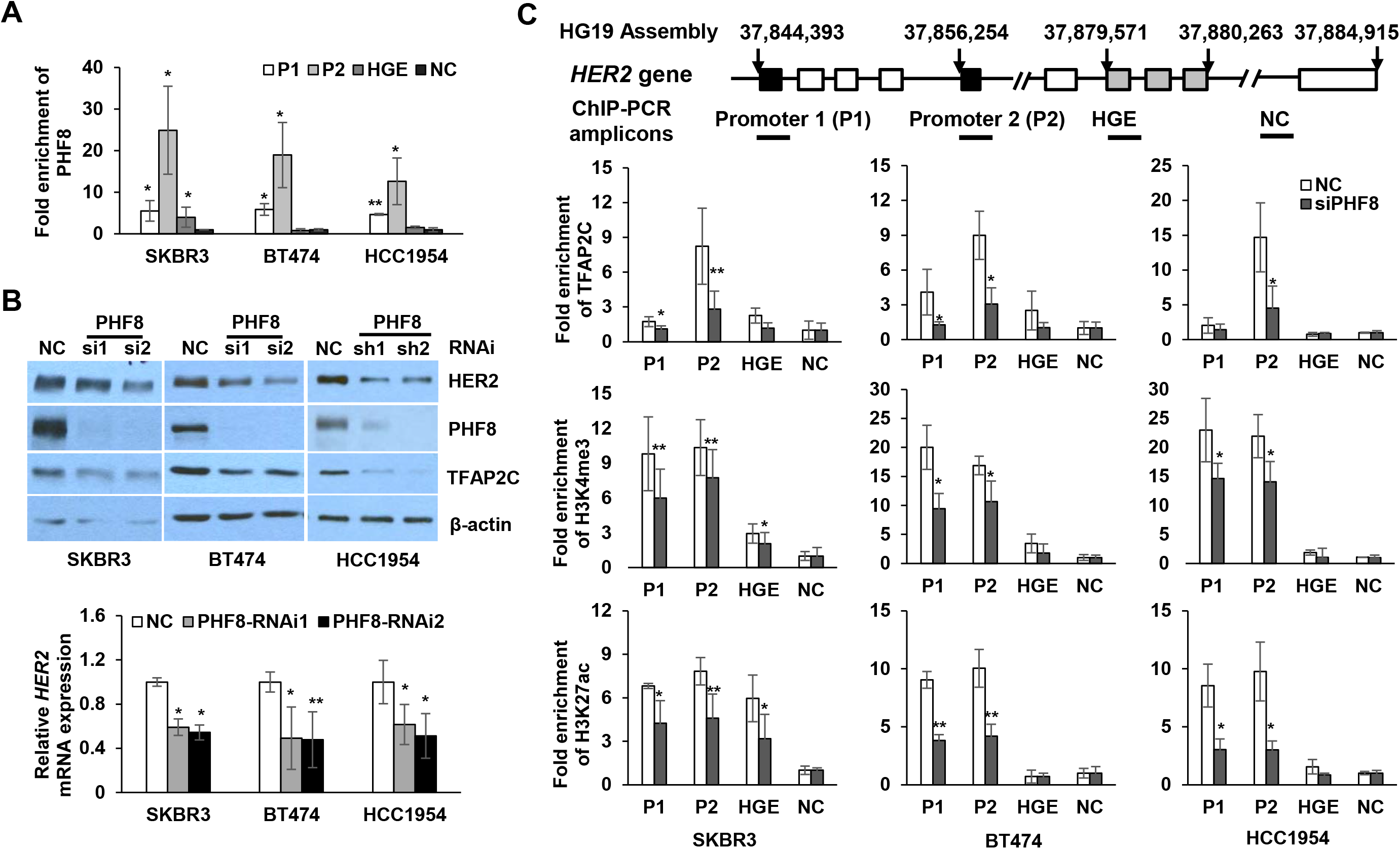
PHF8 plays a transcriptional coactivator role in *HER2* expression. **A.** PHF8 chromatin immunoprecipitation (ChIP) on *HER2* gene in HER2+ cells. Enrichment of PHF8 on *HER2* promoter 1 (P1), promoter 2 (P2), *HER2* gene body enhancer (HGE) and a region clear of modification (NC) were analyzed by ChIP-qPCR. Relative enrichment represents average fold enrichment in PHF8 ChIP vs. input, normalized to NC region. Values are represented by means ±SD from four independent ChIP experiments. * *p*<=0.05, **: *p*<=0.01. **B**. HER2 protein and mRNA levels were assessed by western blotting and RT-PCR in the indicated cells with and without PHF8 knockdown. NC: control scrambled siRNA or shRNA; PHF8-si1 and si2 are PHF8 siRNAs; PHF8-sh1 and sh2 are PHF8 shRNAs. *HER2* mRNA levels were normalized over *RPL13A*. **C**. The enrichments of H3K27ac, H3K4me3 and TFAP2C on *HER2* gene in HER2+ cells with or without PHF8 knockdown were assessed by ChIP-qPCR. Student’s t-test was performed. * *p*<=0.05, **: *p*<=0.01.

### PHF8 has dominant coactivator function downstream of HER2 signaling

Although, PHF8 directly participates in the transcriptional regulation of *HER2, PHF8* knockdown only reduced about 30% of *HER2* mRNA levels (Figure 2B). Thus, we aim to further dissect the genome-wide impact of PHF8 on HER2-regulated genes. MCF10A cells have been extensively used to study HER2 function (Bollig-Fischer et al, 2010; Kim et al, 2009; Yong et al, 2010). Thus, we established double-stable cell lines using MCF10A cells that stably overexpress HER2 and doxycycline-inducible control or *PHF8* shRNAs. As HER2 can induce genomic instability (Burrell et al, 2010), we use early passages of these cell lines to maintain their isogenic status. Consistent with previous reports (Ingthorsson et al, 2015; Kim et al, 2009), HER2 overexpression induces proliferation, AKT phosphorylation (p-AKT) and EMT markers (N-Cadherin (CDH2), ZEB1) (Figure 3A and B). Notably, silencing of PHF8 attenuated these inductions (one shRNA is shown, but both shRNAs had the same phenotype) (Figure 3A and B). Interestingly, *PHF8* knockdown also downregulated the overexpressed HER2, implicating that PHF8 may indirectly regulate HER2.

**Figure 3.**
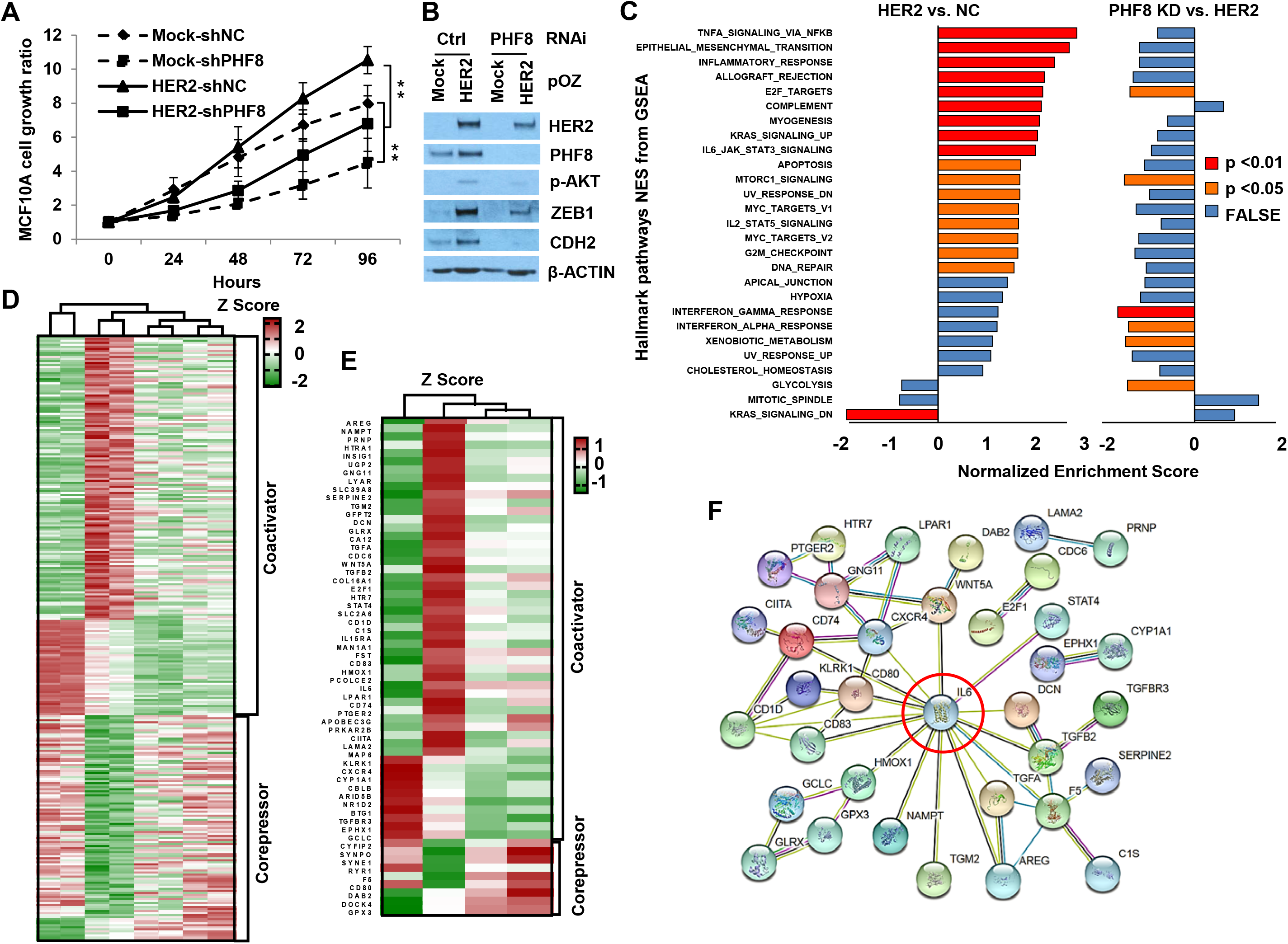
PHF8 plays transcriptional regulatory roles on HER2-induced genes. **A.** The proliferation of MCF10A cells with either overexpression of HER2 or PHF8 knockdown or both was assessed by MTT assay. The data are interpreted as means ± SD from three independent experiments. \*\**p*<=0.01. **B.** HER2, PHF8 and other indicated proteins from the MCF10A cell lines as in A were examined by western blotting. The data are representative from three independent experiments. For both **A** and **B**, Mock: overexpression control; HER2: HER2 overexpression; shNC: scrambled shRNA; shPHF8: PHF8 shRNAs. **C.** PHF8 knockdown counteracts most of the pathways induced by HER2 overexpression. Gene-Set Enrichment Analysis (GSEA) of Hallmark pathways of the genes significantly regulated by HER2 overexpression compared with PHF8 knockdown. Nominal p-values were shown by colors as indicated; NES (normalized enrichment scores) were used as x-axis. **D.** Heat map of RNA sequencing Z-score results of 298 PHF8 differentially regulated genes (DRG) that are significantly regulated by HER2 overexpression. **E.** Heat map of Z-scores of 60 PHF8 DRGs contributing most to enriched pathways. **F.** Protein-protein association networks of 60 PHF8 DRGs were analyzed by STRING. Edges represent protein-protein associations such as interactions, gene neighborhood, co-expression and co-occurrence.

We next we carried out RNA-seq on these cell lines expressing mock/control shRNA, HER2/control shRNA and HER2/PHF8shRNA 1 and 2. The RNA-seq data were analyzed with kallisto pseudo alignment (Bray et al, 2016) and the differential expression was determined using the sleuth R package (Pimentel et al, 2017). 838 upregulated and 536 downregulated genes by HER2 overexpression were obtained using cutoff of actual fold change of 1.5 (FC >=1.5 or <=-1.5) and adjusted *p* value of less than 0.05 (Supplemental Table 2). Gene Set Enrichment Analysis (GSEA) (Subramanian et al, 2005) analysis with Hallmark pathways revealed that HER2-regulated genes are significantly enriched in 18 pathways such as TNFα signaling, EMT transition, inflammatory response and mTOR (Figure 3C and Supplemental Table 5), consistent with previous reports with similar set up (Dong et al, 2017; Liu et al, 2018b; Pradeep et al, 2012). PHF8 knockdown attenuated most of the pathways induced by HER2 overexpression (Figure 3C and Supplemental Table 6). Importantly, the pathways of E2F targets, mTOR and interferon responses elevated by HER2 overexpression are significantly counteracted by PHF8 loss-of-function (Figure 3C). These data strongly suggest a general coactivator functions of PHF8 in HER2-induced transcriptome.

We next defined 298 PHF8-differentially regulated genes (DRG) subtracted from HER2-regulated genes with following criteria: the same trend of regulation by two PHF8 shRNAs; at least one shRNA shows more than 30% of the regulation and the *p* value is less than 0.05. The 30% cut off is based on the general regulatory function of PHF8 from several published data (Liu et al, 2010; Qi et al, 2010; Wang et al, 2016). These 298 DRGs were clustered into two groups based on the transcriptional functions of PHF8: coactivator (PHF8 knockdown attenuated the genes upregulated by HER2 or enhanced the genes downregulated by HER2); corepressor (PHF8 knockdown counteracted the genes downregulated by HER2 or enhanced the genes upregulated by HER2) (Figure 3D and Supplemental Table 3 and 4). These analyses led to a general conclusion: the transcriptional coactivator function of PHF8 is dominant over its corepressor function. These genes were further analyzed for GO biological processes through Enrichr and show that PHF8 coactivator genes (upregulated by HER2) are enriched in cell proliferation and cytokine production, whereas, PHF8 corepressor genes (downregulated by HER2) are enriched in axon and neuron regeneration (Supplemental Figure 5).

To further investigate the biological impact of PHF8 on HER2-regulated genes, the 298 DRGs were subtracted from the genes contributed to significantly enriched pathways regulated by either HER2 overexpression or PHF8 knockdown and led to a 60 gene signature (Figure 3E and Supplemental Table 7). These genes further demonstrate the dominant coactivator functions of PHF8 downstream of HER2 signaling. Analysis of protein-protein association networks of these 60 genes by STRING (Szklarczyk et al, 2019) revealed that Interleukin-6 (IL-6) is the hub connecting to other molecules (Figure 3F). In fact, IL-6 contributes to 9 pathways such as interferon response, TNFα signaling, EMT, and inflammation (Supplemental Table 7). These data suggest that PHF8 may contribute to the oncogenic functions of HER2 through IL-6.

### PHF8 upregulates IL-6 and contributes to trastuzumab resistance

We next sought to pursue the regulation of IL-6 by PHF8 and how this regulation contributes to the resistance to anti-HER2 drugs based the following facts: the regulation of *IL-6* by HER2 overexpression ranks higher (Supplemental Table 7); the central position of IL-6 in the protein-protein association network of PHF8 DRGs in context of HER2 (Figure 3F); the functional importance of IL-6 in drug resistance (Conze et al, 2001; Ghandadi & Sahebkar, 2016) and in HER2 signaling (Chung et al, 2014). First, we confirmed the regulation of *IL-6* by HER2 and PHF8 in the MCF10A cell lines (Figure 4A). Such transcriptional regulation of *IL-6* by PHF8 was further confirmed in HCC1954 and BT474 lapatinib-resistant (-R) (Stuhlmiller et al, 2015) cells (Figure 4B), which possess higher *IL-6* mRNA and protein levels compared with other cell lines tested (Supplemental Figure 6). We next carried out human cytokine antibody array (RayBio C-Series Human Cytokine Antibody Array 5) and obtained consistent results: PHF8 knockdown attenuated the upregulation of IL-6 by HER2 overexpression in MCF10A cells (Figure 4C) and downregulated IL-6 in HCC1954 cells (Figure 4D). The regulation of IL-6 by PHF8 in HCC1954 cells was further validated by ELISA assay (Figure 4E). Notably, ANG (Angiogenin) and CCL20 are also regulated by HER2 and PHF8 in the similar pattern as IL-6 in MCF10A cells (Figure 4C). However, such regulation by PHF8 was not repeatable in HCC1954 cells (Figure 4D).

**Figure 4.**
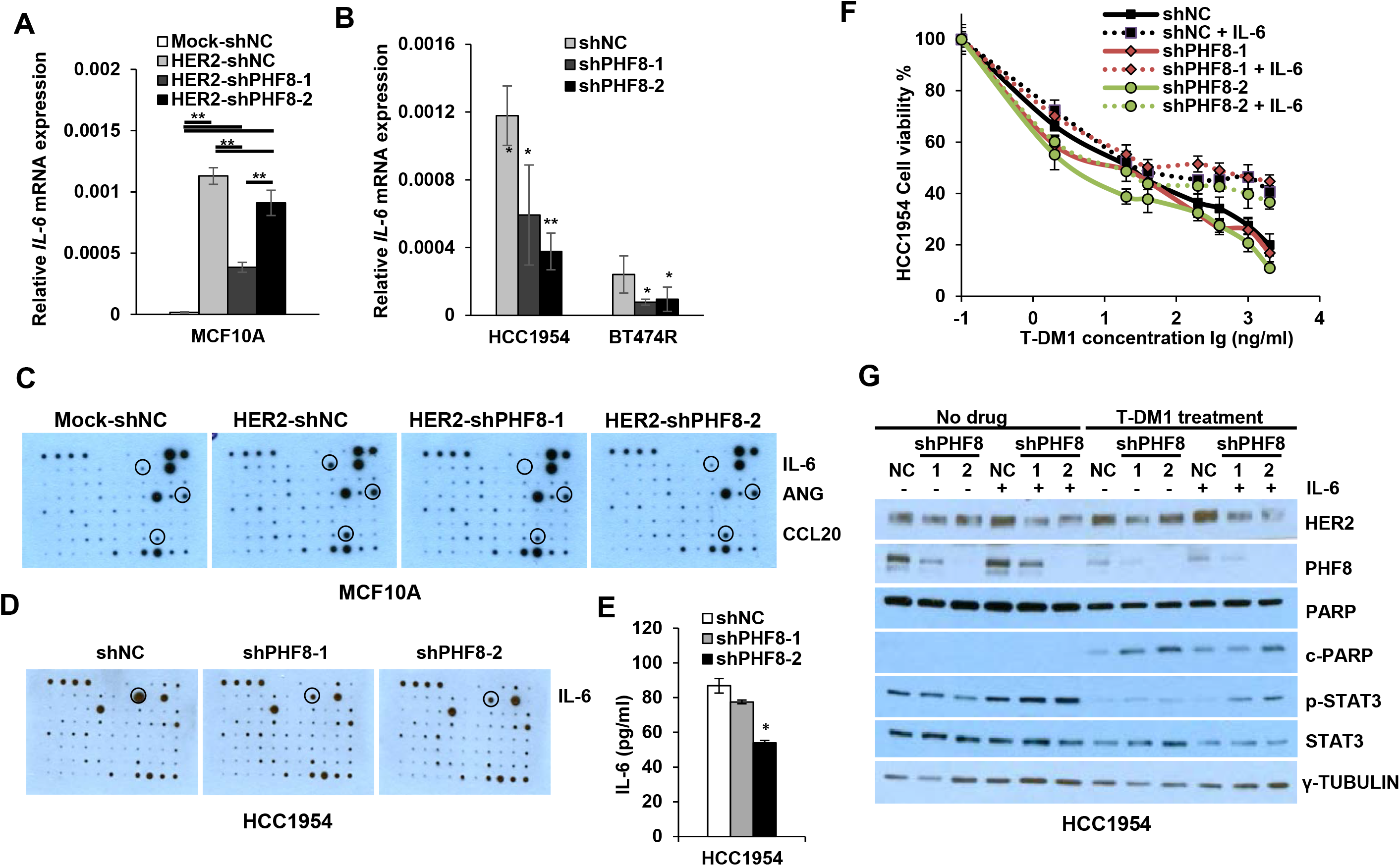
PHF8 upregulates IL-6 and contributes to trastuzumab resistance. **A** and **B**. Relative expression of *IL-6* was assessed in the indicated cells by RT-qPCR. *RPL13A* was served as loading control. \**p*<=0.05; \*\**p*<=0.01. **C** and **D**. Human Cytokine Antibody Array blots probed with the media from the indicated cells. Duplicated blotting was performed and the regulated cytokines are marked. E. Secreted IL-6 from HCC1954 cells with or without PHF8 knockdown was determined by Quantikine ELISA. \**p*<=0.05; \*\**p*<=0.01. **F.** Cell viability of HCC1954 cells with or without PHF8 knockdown, and addition of IL-6 as indicated was examined by MTT assay. Cells were treated doxycycline for 72 hours, reseeded in 96-well plates. IL-6 (100 ng/mL) was added to the cells 24 hours prior to the T-DM1 treatments (48 hours) (n=5). **G.** The levels of HER2, PHF8, PARP/c-PARP (cleaved PARP) and STAT3/P-STAT3 were examined by western blotting in HCC1954 cells treated as in **F**, γ-Tubulin was used as loading control. The data are representative of three independent experiments.

As HCC1954 cells possess *de novo* trastuzumab resistance (Sahin et al, 2009) we next tested if PHF8 and PHF8-IL-6 axis contribute to such resistance. The IC_50_ of Trastuzumab (T-DM1) in HCC1954 cells with control shRNA is 92.44 ng/ml as determined by MTT assay (Figure 4F). PHF8 knockdown by two shRNAs reduced the IC_50_ to 33.72 ng/ml and 29.68 ng/ml, respectively (Figure 4F), suggesting a positive role of PHF8 in the trastuzumab resistance of HCC1954 cells. When exposing to IL-6 (dotted lines in Figure 4F), the HCC1954 cells with control shRNA did not die following a linear pattern within increased concentrations of T-DM1. IC_50_ value can be roughly calculated as 106.43 ng/ml (Figure 4F). However, IL-6 does elevated the IC_50_ to 104.57 ng/ml and 64.47 ng/ml in PHF8-knockdown cells, supporting the contribution of PHF8-IL6 axis to the trastuzumab resistance of HCC1954 cells. Next, western blotting was performed on these cells with and without treatment of T-DM1 at 2 ng/ml (Figure 4G). At static state, PHF8 knockdown slightly reduced the activated STAT3 (p-STAT3) and addition of IL-6 restored the p-STAT3 levels (Figure 4G), supporting the role of PHF8 in regulating IL-6 signaling. T-DM1 treatment in the control cells significantly reduced p-STAT3 and induced apoptosis reflected by cleaved PARP (c-PARP) (Figure 4G). PHF8 knockdown increased the T-DM1-induced apoptosis and addition of IL-6 counteracts the apoptotic induction (Figure 4G), supporting the contribution of PHF8-IL-6 axis to the trastuzumab resistance of HCC1954 cells.

### *Phf8* contributes to Her2-driven breast tumor development *in vivo*

We previously revealed the functional requirement of PHF8 in the anchorage-independent growth of HER2+ breast cancer cells (Shao et al, 2017). Our current data support the synergistic interplay between PHF8 and HER2. Thus, we sought to further study the role of PHF8 in HER2-driving tumor development *in vivo: Phf8* knock out mouse model with *Her2* overexpression driven by mouse mammary tumor virus *(MMTV)* long terminal repeat (LTR) promoter. The *Phf8^flox/flox^* allele was established by flanking exon 8 with two *loxp* cassettes (Figure 5A). Exon 8 of *Phf8* encodes amino acids 261-316, part of the C-terminal JmjC domain containing a 2-oxoglutarate (2-OG) binding residue (K264) (Figure 5A). Deletion of this region causes truncation of Phf8 and abolishes its demethylase activity. We used *MMTV-Cre* to drive *Phf8* knockout (KO) in mammary epithelial cells. Genotyping demonstrated successful insertion of the *loxp* cassettes (Figure 5B). Next, *MMTV-Her2, MMTV-Cre* and *Phf8^flox/flox^* mice were intercrossed to generate wild type PHF8 (referred as WT) mice: *MMTV-Her2/MMTV-Cre, MMTV-Her2/Phf8^flox/flox^* and the PHF8 KO mice: *MMTV-Her2/MMTV-Cre/Phf8^flox/flox^*. Notably, due to the X chromosome localization of *Phf8*, portion of *MMTV-HER2/MMTV-Cre/Phf8^flox/wt^* can possess total loss of Phf8 due to X chromosome inactivation, an example of such case is shown in Figure 5C (lane 2). 42 WT mice and 44 *Phf8* KO mice all with *MMTV-Her2* background were collected. All these mice were genotyped (tail) (Figure 5B shows selected mice) and PHF8 protein levels from all tumor samples were verified by western blotting (Figure 5C, lanes 1,2,5 and 6 versus lanes 3-4). spontaneous mammary gland tumors developed from 185.8 ± 42.8 (mean ± SD) and 180 ± 39.2 days in WT and *Phf8* KO mice, respectively, no significance was obtained (Figure 5D). However, the tumor weight (2.64 ± 1.26 g) from *Phf8* KO mice was significantly decreased compared with that from WT mice (3.22 ± 1.40 g) (Figure 5E). Moreover, tumor ratio (relative tumor weight percentage of total body weight) from *Phf8* KO mice was significantly reduced to 6.97% from 9.31% from WT mice (Figure 5F). These data revealed that Phf8 plays critical roles in tumor growth rather than tumor initiation. This conclusion was further supported by significantly reduced proliferative index (KI67 IHC staining) in *Phf8* KO mice (n=7) versus the WT mice (n=6) (12.47 ± 4.56 % vs. 3.57 ± 2.18%; *p* = 0.002).

**Figure 5.**
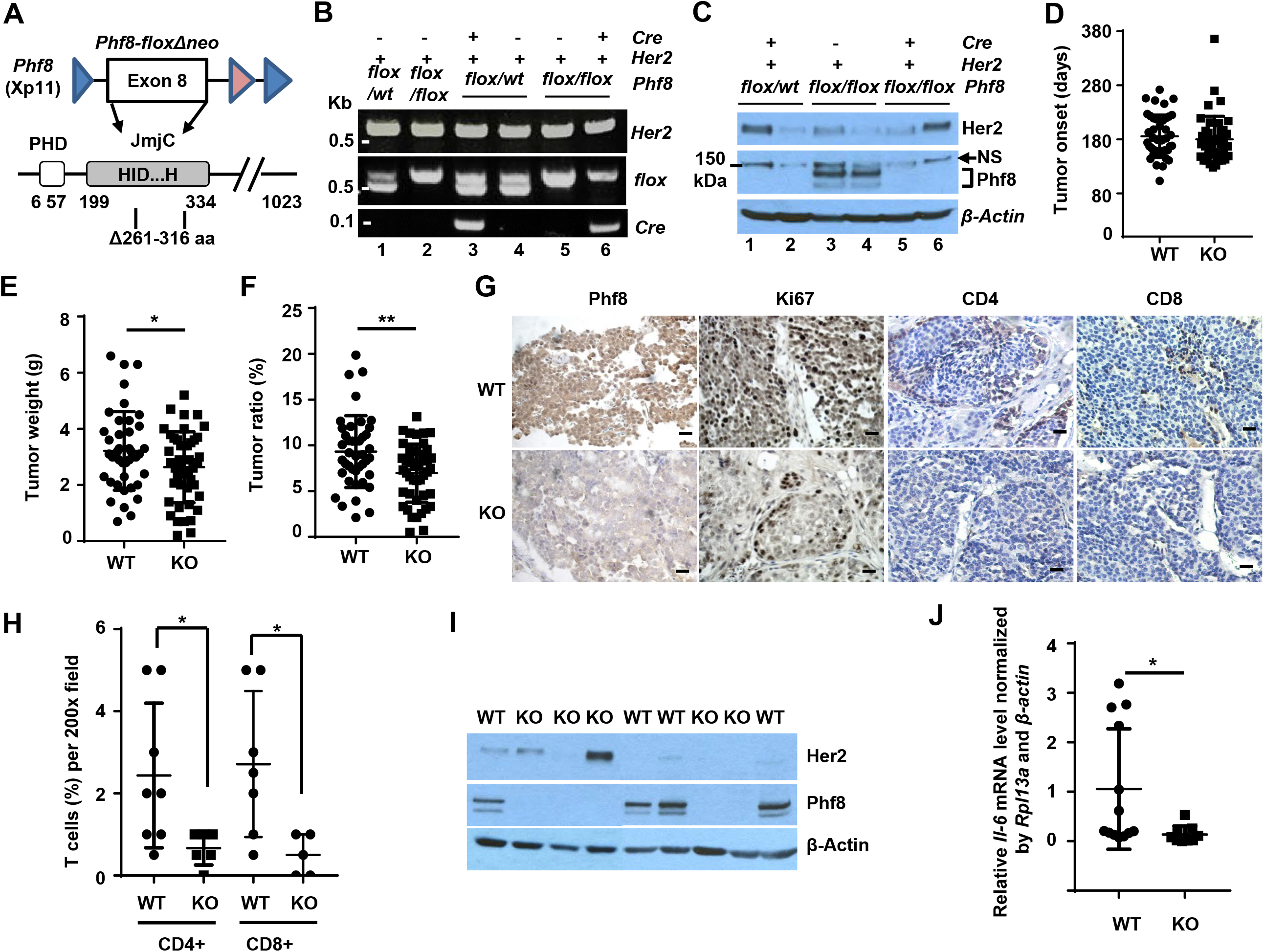
Functional impact of PHF8 on HER2 tumorigenesis and T cell infiltration *in vivo*. **A.** Schematic illustration of the floxed locus in mouse *Phf8* gene and the corresponding coding amino acids. **B.** Genotyping of *Her2, flox* and *Cre* using tail DNA from the indicated mouse. kbp: Kilo base pair. **C.** Phf8 and Her2 protein levels from mice with the indicated genotypes were assessed by western blotting. **D-F.** Tumor onset date (**D**), tumor weight (**E**) and tumor ratio (**F**) from WT and Phf8 KO mice. Each point represents an individual animal. Student’s t-test was used to calculate the significance between KO and WT groups. N=86. **G.** IHC staining of Phf8, Ki67, CD4 and CD8 in PFA-fixed, paraffin-embedded tumor tissues from *Phf8* KO and WT tumor section (magnification, ×200). Bar: 10 μm. **H**. Comparison of infiltrating T cells in tumors from WT and *Phf8* KO mice. The percentage of both intraepithelial and stromal tumor-infiltrating CD8+ and CD4+ T cells was quantified per 200x field. **I**. HER2 and Phf8 levels in the primary tumor cell lines from *Phf8* KO and WT mice were assessed by western blotting of. **J**. mRNA levels of *Il-6* were examined from the primary tumor cell lines by RT-qPCR. *Il-6* expression was normalized by both *Rpl13a* and *β-actin*. \**p*<=0.05.

Given the role of PHF8 in regulation of IL-6 and the well-established IL-6-immunoresponse network in tumor growth (Fisher et al, 2014; Mauer et al, 2015; Tsukamoto et al, 2015), we next analyzed infiltrating T cells. Compared with wild spread T cells from WT mice (N>=7), the tumors from *Phf8* KO mice (N>=5) show nearly no intratumoral T cells but few peripheral to the tumor mass and in collagen bundles (Figure 5G). IHC staining for CD4 and CD8 further revealed significant reduction of intratumoral and peri-tumoral infiltrating T cells in Phf8 KO mice compared with WT mice (Figure 5H). These data consistent with studies indicating IL-6 have dual function in tumor microenvironment: it can promote inflammation-induced CD8+ T cell trafficking in tumors (Fisher et al, 2011) at the same time include stromal cells like CD4+ regulatory T cells (Treg) supporting tumorigenesis. Next, we aim to examine if the regulation of Il-6 by Phf8 is conserved. Due to mixed cell population in the tumors, we established 13 WT and 9 KO primary tumor cell lines from WT (n=8) and KO (n=6) mice, respectively. Her2 and Phf8 levels were verified by western blotting (Figure 5I). Importantly, the *Il-6* mRNA levels in cells originated from KO mice are significantly lower than that from the WT mice (Figure 5J), consistent with our data from cell lines. Taken together, these data implicate that PHF8-IL-6 axis may mediate the T cell infiltration in HER2-driving tumor development.

## Discussion

Therapy resistance to anti-HER2 drugs such as to Lapatinib or Trastuzumab remains a hurdle in successful therapy of HER2+ breast cancers (Roskoski, 2014). Thus, to identify novel therapeutic target is critical. Epigenetic factors that can be inhibited with specific inhibitors may serve for this purpose. With *in vitro* and *in vivo* approaches, this study demonstrates the synergistic interplay between histone demethylase PHF8 and HER2 and the oncogenic functions of PHF8 in HER2-driven tumor development, conferring therapeutic significance targeting PHF8 in HER2+ breast cancers.

Oncogenic functions of PHF8 have been recognized in various types of cancer (Bjorkman et al, 2012; Maina et al, 2016; Shen et al, 2014; Sun et al, 2013; Zhou et al, 2018). We (Shao et al, 2017) and Wang Q et al (Wang et al, 2016) discovered such functions of PHF8 in breast cancers and reported the significant increase of PHF8 mRNA levels in several subtypes of breast cancers. However, our previous (Shao et al, 2017) and current approaches with updated dataset containing increased samples (291 normal and 1085 breast cancer samples) did not achieve a significant upregulation of PHF8 mRNA levels in HER2+ breast cancers. In contrary, our IHC data from a pool of over 500 samples revealed significant increase of PHF8 protein levels in all subtypes of breast cancers. Despite of the fact that the mRNA data are not the same as our breast cancer array samples, the discrepant mRNA and protein levels of PHF8 in breast cancers re-emphasize the roles of the post-transcriptional (Shao et al, 2017) and post-translational (Wang et al, 2016) regulations of PHF8. Although, our study showed the transcriptional regulation of *PHF8* by exogenous HER2, the c-MYC-miR-22-PHF8 regulatory axis (Shao et al, 2017) may still apply to the regulation of PHF8 by HER2 signaling due the following reasons: 1) MYC has long been recognized as a key player in HER2-mediated cancerous transformation (Hynes & Lane, 2001; Nair et al, 2014); 2) miR-22 expression is lowered in HER2+ breast cancers (Mattie et al, 2006); 3) our previous study showed that miR-22 mimics downregulate PHF8 protein level in SKBR3 cells (Shao et al, 2017). Beyond miR-22, let-7 also targets PHF8, adding a possible HER2-let-7-PHF8 axis, because: 1) let-7 levels are lower in HER2+ breast cancers (Mattie et al, 2006); 2) HER2 can repress let-7 expression through ERK signaling (Liu et al, 2015) by activating LIN28, which inhibits let-7 family biogenesis (Paroo et al, 2009); 3) HER2 can also inhibit let-7 through a different pathway involving AKT-MYC (Chang et al, 2009). Taken together, HER2 may regulate PHF8 through multiple mechanisms. Further studies are needed to dissect the HER2-microRNAs-PHF8 axis.

Through RNA-seq and pathway analyses, we found that the transcriptional coactivator role of PHF8 downstream of HER2 signaling is dominant over its corepressor role, consistent with most of the studies (Liu et al, 2010; Qi et al, 2010; Shao et al, 2017; Wang et al, 2016). As higher transcription rate of *HER2* per gene copy was observed in HER2-amplified breast cancer cells (Bofin et al, 2004; Kraus et al, 1987; Mungamuri et al, 2013), adding PHF8 to MLL complex and H3K9 acetyltransferase (Mungamuri et al, 2013) helps to decipher the epigenetic regulatory machinery of *HER2* gene. Toward the demethylation substrates of PHF8, although, we reasoned that the constitutively active *HER2* gene may have lower H3K9me2 levels, it is still possible that PHF8 plays a role to sustain the low occupancy of H3K9me2 on HER2 genomic region. Furthermore, *HER2* gene can be transcriptionally upregulated by tamoxifen, an ER antagonist, in ER+ breast cancers and by radiation therapy triple-negative breast cancers, hence activated HER2 signaling contributes to resistance of endocrine or radio-therapy (Cao et al, 2009; Duru et al, 2012; Hurtado et al, 2008). It would be very interesting to learn if PHF8 plays a transcriptional coactivator role in breast cancer cells with *HER2* overexpression in addition to *HER2* amplification.

Beyond *HER2*, the HER2 signaling-specific DRGs regulated by PHF8 are enriched in positive regulating cell proliferation followed by chemokine/cytokine production pathways, implicating the role of PHF8 in the immuno-response of tumors in addition to its well-known functions in cell cycle regulation. IL-6, CD74, TGFB2, WNT5A and HMOX1 are among the major components for the cytokine-related pathways. IL-6 signaling is considered as a malevolent player (Fisher et al, 2014). However, IL-6 also has obvious tumor-inhibitory effects as it influences tissue recruitment of T cells (Hunter & Jones, 2015) and works as a key player in the activation, proliferation and surviving of lymphocytes during active immune responses promoting anti-tumor adaptive immunity (Fisher et al, 2014). TGF-β and WNT signaling pathways play key roles in EMT and cancer stem cells gaining cancer metastasis and resistance to therapies, meanwhile, they can also have the immuno-repressive functions (Esquivel-Velazquez et al, 2015). The coactivator functions of PHF8 on these cytokines may lead to controversial outputs, however, the reduction of infiltrating T cells in the tumors from Phf8 KO mice demonstrates the positive role of PHF8 in tumor T cell infiltration. Thus, PHF8 may play important role in immuno response by change tumor microenvironment and influence T cell trafficking to tumor sites by regulating cytokine production.

In addition to the dominant coactivator function of PHF8, studies also show that PHF8 has corepressor function, i.e. PHF8 is phosphorylated by ERK2 upon IFNγ treatment and is evicted from repressive promoters, where PHF8 forms a complex with HDAC1 and SIN3A at the static state (Asensio-Juan et al, 2017). This mechanism may also apply to the context of HER2 because ERK can be activated by the activated HER2 signaling (Baselga & Swain, 2009; Roskoski, 2014). Genome-wide PHF8 ChIP-seq in the MCF10A cell system is sought to decipher how PHF8 plays its corepressor function on the genes identified in this study.

## Materials and Methods

### Cell lines and treatments

All cell lines used in this study were obtained from the American Type Culture Collection (ATCC) (Rockville, MD, USA). MCF10A cells were cultured in DMEM-F12 supplemented with 20 ng/ml Epidermal Growth Factor (EGF) (Sigma), 100 ng/ml cholera toxin (Sigma), 10 g/ml insulin (Sigma), 500 ng/ml hydrocortisone (Sigma), and 5% horse serum. HCC1954, SKBR3 and BT474 cells were cultured in RPMI1640 medium containing 10% FBS. HEK293T cells were grown in Dulbecco’s Modified Eagle Medium (DMEM) containing 10% FBS (Gibco). All these cell lines were maintained in the specified medium supplemented with 1× Penicillin— Streptomycin (Gibco) and incubated in 5% CO_2_ at 37°C. MCF7-HER2 and MCF10A-HER2 (overexpressing HER2) were established using pOZ retroviral system as described previously (Qi et al, 2010). Cell lines stably expressing PHF8 shRNAs were established as described before (Shao et al, 2017). Briefly, HEK293T cells in a 10 cm dish with about 60% confluence were transfected with 8 μg lentiviral construct and helper plasmids using Lipofectamine 2000 (Life Technologies). 24 hours after transfection, culture media were changed and supernatants were collected 48 hours later. The cells were infected with the virus and selected by puromycin (1-2.5 μg/mL) for 10 days. Knockdown efficiency was verified by quantitative RT-PCR. Two different shRNAs per target gene were tested to reduce off-target effects. T-DM1 (Kadcyla, Genentech) was used to treat cells for 24 hours. Dimethyl sulfoxide (equal volume to that of treated cells) was added to culture media of the control cells. Human Cytokine Antibody Array blots probed with the cell cultured media. Serum free media were added after doxycycline induction for 72 hours of each cell line and collected after 24 hours. Recombinant human IL-6 was purchased from PeproTech (Rocky Hill, NJ). Enzyme-linked immunosorbent array (ELISA), 10,000 cells were seeded in 6-well plates with complete medium of each cell line with FBS for 24 hours. The medium was then replaced to serum-free medium for another 24 hours, and the supernatant was tested using the Quantikine human IL-6 (sensitivity < 5 pg/mL) ELISA kit (R&D Systems, Inc., Minneapolis, MN) according to manufacturer’s recommended conditions. Cells were induced with doxycycline for 72 hours first before serum-free medium replacement, and the cytokine in the media was analyzed by ELISA.

### RNA-seq analysis

RNA sequencing (RNA-seq) was performed on duplicated RNA samples from MCF10A cells overexpressing empty vector (Mock), or HER2, in combination with control scrambled shRNA or two PHF8 shRNAs: mock/shNC, HER2/shNC, HER2/PHF8shRNA1 and HER2/PHF8shRNA2. Expression values were calculated as FPKM (fragment per kilobase of exon per million of mapped fragments) and were used to determine differential expression of mRNAs in four groups of samples. Transcripts were called expressed if FPKM values in overexpression of HER2 with shRNA control samples were ≥1.0 for mRNAs. The mean expression level and differences in expression between four groups were calculated, and from these statistically significant differences in expression between each group was determined using a paired *t* test. Fold change (HER2/shNC versus mock/shNC) was calculated to identify differentially expressed transcripts. Transcripts were deemed differentially regulated by HER2 overexpression with criteria of fold change ≥1.5 or ≤0.5 and *p* value less than or equal to 0.05. Additionally, transcripts were deemed as PHF8 conserved regulated if mean fold change of HER2/ PHF8shRNA1 or 2 versus HER2/ctlshRNA was ≥1.3 or ≥0.7 (*p* ≤ 0.05) and regulation by two shRNAs were of same trend. Differentially expressed mRNAs that are significantly regulated by PHF8 knockdown are represented by heatmaps by Graphpad prism 7, and Z scores were scaled by row using standard Z score calculation of log 10 absolute FPKM values.

### ChIP and ChIP-qPCR

Chromatin immunoprecipitation (ChIP) was performed as described previously (Shao et al, 2017). Briefly, formaldehyde crosslinked cells were lysed and sonicated to shear the DNA. The sonicated DNA-Protein complexes were immunoprecipitated with the following antibodies: control IgG (A01008, GenScript), anti-TFAP2C (sc-12762, Santa Cruz), anti-PHF8 (ab36068, Abcam), anti-H3K4me3 (ab8580, Abcam), anti-H3K27ac (ab4729, Abcam). The immuno complexes were collected using protein A/G agarose beads. The eluted DNA and 10% of respective input DNA were reverse cross-linked at 65°C overnight and used for the qPCR using SYBR Green qPCR mix and a CFX96 instrument (BioRad).

### Cell Proliferation Assay (MTT)

Cells were seeded at a density of 3×10^3^ cells/well in a 96-well plate with outer wells left empty for addition of PBS. After 24 hours of culture, the media was changed and vehicle, drugs, or IL-6 (100 ng/mL) (Block et al, 2012) were added. The cells were incubated with inhibitors or drugs for the time specified; then 0.5 mg/mL MTT dye was added and the cells were incubated for an additional 4 hr. Formazan crystals were dissolved in dimethyl sulfoxide (DMSO) for 15 min and the plates were read spectrophotometrically at 590 nm with a reference of 650 nm. Each assay was performed at least three times with five wells replication.

### Mouse works

All mouse work was performed under protocols approved by the Institutional Animal Care and Use Committee (IACUC) at The University of Iowa. *Phf8* knockout mice were established by Dr. Yang Shi’s lab at Harvard Medical School. MMTV-Her2 mice were provided by Dr. Weizhou Zhang. All mice have been fully backcrossed to FVB/N mice (from the Jackson Laboratory) for 8 generations. Her2-driven mammary tumors were monitored every 3 days. At experimental endpoint when largest tumor reaches 2 cm in diameter, animals were sacrificed, and all mammary tumors were removed and weighed. After the mice were sacrificed, tumor weight was directly measured, and tumor ratio was calculated as percentage of body weight. Tumors were fixed, embedded in paraffin, and serially sectioned at a thickness of 6-8 μm, and IHC staining was performed as described previously (Shao et al, 2017).

### Statistical analysis

GraphPad Prism software (v7) was used to conduct statistical analysis. Results are interpreted as mean ± SD. Differences between experimental groups were compared using an unpaired two-tailed Student’s *t*-test (for two conditions). A \**p* value <=0.05 was considered statistically significant. ** *p* value <=0.01 was considered highly significant.

### Oligonucleotides

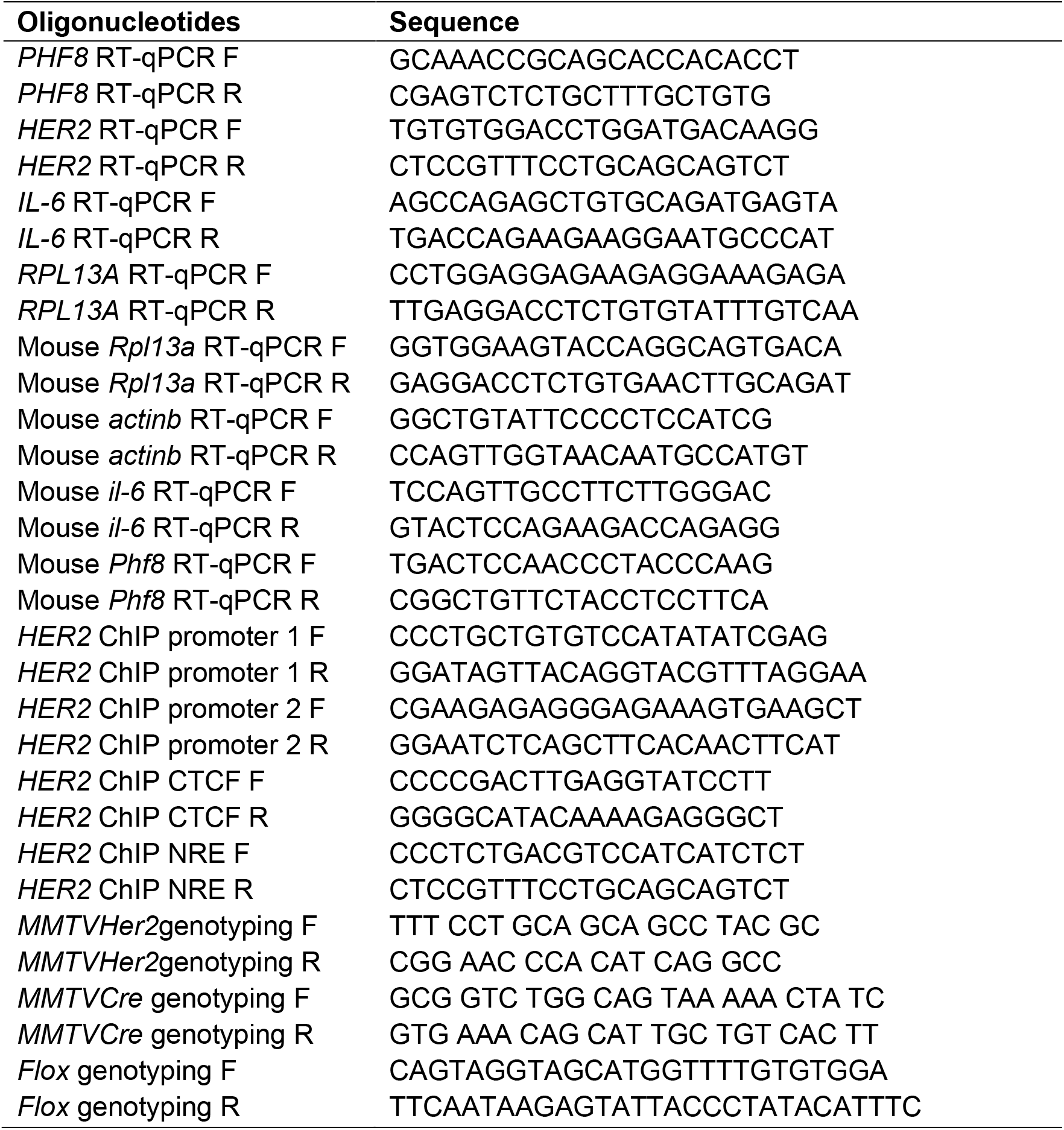
All oligonucleotides were synthesized by IDT and the sequences are shown below.

## Acknowledgments

We thank Dr. Yang Shi and Dr. Hui-Jun Lim for providing *Phf8* knockout mice. We also thank Dr. Songhai Chen and Drs. Sonia Sugg, Amani Bashir for technical assistance on the mouse work and the IHC data analysis, Dr. Brad Amendt and his lab for helpful discussions. This work was supported by lab start-up funds to H.H.Q from the Department of Anatomy and Cell Biology, the Carver College of Medicine, University of Iowa; Carver Trust Young Investigator Award (01-224 to H.H.Q) from the Roy J. Carver Charitable Trust; a Breast Cancer Research Award (to H.H.Q.) by the Holden Comprehensive Cancer Center at University of Iowa; The NIH grant (P30 CA086862) to the Genomics facility at the University of Iowa. N.B. was supported by NIH M.D./Ph.D. fellowship (F30 CA206255); W.Z was supported by NIH grants CA200673, and CA203834, the V Scholar award, a Breast Cancer Research Award and an Oberley Award (National Cancer Institute Award P30CA086862) from Holden Comprehensive Cancer Center at the University of Iowa.

## Author Contributions

QL and HHQ conceived the concept of the paper and co-wrote the paper. QL carried out most of the experiments. NB carried out the bioinformatic analysis. PS and PKM contributed to the RNA-seq and vector constructions. WZ supervised the mouse work and participated in the manuscript writing.

## The paper explained

1. Novel epigenetic mechanisms by which the histone demethylase PHF8 interplays with HER2 and plays critical roles in HER2-driven tumor development and the resistance to anti-HER2 drugs.
2. PHF8 is elevated in HER2-positive breast cancers and is upregulated by HER2;
3. PHF8 plays coactivator roles in regulating *HER2* expression, HER2-driven epithelial-to-mesenchymal transition (EMT) markers and cytokines;
4. HER2-PHF8-IL-6 regulatory axis was proved, and it contributes to the resistance of Trastuzumab *in vitro* and may play a critical role in the infiltration of T-cells in HER2-driven breast cancers.

## For more information

Not applicable

## Data availability

The RNA-seq data are in the process to submit to GEO database.

